# Deep single-cell immune and signaling profiles predict long term therapy response in chronic myeloid leukemia within hours

**DOI:** 10.1101/2025.11.10.687542

**Authors:** Stein-Erik Gullaksen, Mathias Brevik, Øystein Sefland, Erin Craig, Monica Hellesøy, Ulla Olsson-Strömberg, Jesper Stentoft, Johan Richter, Tobias Gedde-Dahl, Robert Tibshirani, Sylvia Plevritis, Dominic Wolf, Satu Mustjoki, Henrik Hjorth-Hansen, Bjørn Tore Gjertsen

## Abstract

Chronic myeloid leukemia (CML) is effectively treated with small molecule BCR::ABL1 tyrosine kinase inhibitors (TKIs) but like in all cancers, early identification of suboptimal- and non-responders is a challenge. Through mass cytometry analysis of peripheral blood leukocytes, we collected high-dimensional data from *de novo* chronic phase CML patients enrolled in two multicenter clinical trials (clinicaltrials.gov NCT01725204 and NCT01061177). In leukocytes, dasatinib and nilotinib inhibited intracellular signaling within one or three hours after first per oral dose, and each TKI had a unique signaling signature reflecting its kinase specificity profile beyond BCR::ABL1. An immune and signaling profile was constructed for each patient, predicting the treatment response (BCR::ABL1^IS^, %) the first 12 months of treatment. These results show that single cell immune and signaling profiles within hours of first dose can predict 12 months treatment response, anticipating future optimalization of kinase inhibitor treatment within days rather than months.

## Introduction

Response to cancer therapy is monitored by tumor load measurements.(1-3) In chronic myeloid leukemia (CML), a hematopoietic stem cell disease driven by the pathognomonic BCR::ABL1 kinase gene fusion(4-6), response is determined by measuring the residual disease through *BCR::ABL1* mRNA by a standardized RT-qPCR of peripheral blood leukocytes. (7-9) Targeting BCR::ABL1 fusion-protein using tyrosine kinase inhibitors (TKIs) is highly effective, but 30-40% of patients switch their first line choice of TKI treatment.(9-13) The majority switch due to TKI intolerance (2 out of 3), while about 1 out of 3 patients switch due to a lack of response.(11) Treatment free remission is now the therapeutic goal for younger patients with CML, currently only achieved by 20% of newly diagnosed CML patients.(14) Still, most patients require lifelong daily TKI treatment, causing treatment-related side effects and reduced quality of life with significant socioeconomic costs. (15-17)

Dasatinib, a potent sub-nanomolar dual SRC/ABL1 TKI approved for first-line treatment of CML, is readily absorbed in the GI tract and has a mean time to maximum serum concentration of 0.25-1.5 hrs.(18) The SRC family of kinases are efficiently inhibited by dasatinib and *in vitro* experiments have demonstrated that dasatinib reversibly inhibit T cell proliferation and function, in addition to NK cell function.(19-21) However, *in vivo* dasatinib has an immune-modulatory effect with anti-leukemic properties, including expanding subsets of cytotoxic T cells and NK cells.(22-25) Dasatinib induces contraction of stromal cells of the spleen, driving lymphocyte egress and leukocyte mobilization one hour after oral dasatinib intake.(26-28)

Nilotinib, also a second generation TKI approved for first line treatment of CML, has a slower mean time to maximum serum concentration of 3 hours.(29) Nilotinib is a more selective inhibitor of ABL1 compared to dasatinib, but show activity against other kinases including DDR-1/-2, PDGFR-α/β, KIT and CSF1R.(30) The immunological effects of nilotinib on lymphocytes are less explored than dasatinib. At therapeutic concentrations, nilotinib preserves key lymphoid immune functions including T-cell receptor signaling, regulatory T-cell activity, and NK cell cytotoxicity.(31) The primary immune effect of nilotinib has been observed in the myeloid lineages, affecting monocyte function and maturation towards functional dendritic cells (32-34)

The effect of TKIs on the CML leukemic stem cells (LSCs) have been extensively studied since the introduction of imatinib.(35-38) The LSCs are intrinsically resistant to TKI therapy, likely due to their quiescent nature, causing suboptimal response during treatment and relapse after TKI cessation.(39-42) Treatment response is shown to correlate with the total level of phosphorylated tyrosine measured in imatinib-treated CD34^+^ cells.(43) Similarly, *in vivo* inhibition of BCR::ABL1 kinase activity in leukocytes after 7 days of imatinib and nilotinib treatment predict response to treatment.(44) However, few studies have investigated response and the *in vivo* effect of TKIs on the ABL1 signaling pathway across myeloid and lymphoid cells using single cell technologies(45).

We have previously shown the early and direct effects of TKI therapy on both immune and leukemic cells from patients with CML.(39, 46-48) High dimensional mass cytometry analysis suggested that single cell immune and signaling profiling may distinguish optimal responders from sub-optimal responders.(46, 48) Therefore, we hypothesize that it is possible to generate response prediction already hours after the first dose of TKI. We tested this hypothesis in single peripheral blood cells (PB) from two clinical trials of *de novo* chronic phase CML. Analysis of PB collected before and within hours after starting dasatinib or nilotinib therapy demonstrated that the first TKI dose inhibited intracellular signal transduction. Furthermore, single cell profiles of PB collected at baseline and one hour after the first TKI dose predicted response at one year of TKI treatment.

## Results

### Experimental design and mass cytometry derived absolute counts

To investigate the immediate effects of dasatinib (100 mg once daily, NordCML007; NCT01725204) and nilotinib (300 mg twice daily, ENEST1st; NCT01725204) on immune cells, we collected up to four longitudinal peripheral blood samples (PB, n = 229) from each patient with *de novo* chronic phase CML (n=81) (**Figure–1A**). The leukocytes were fixed immediately following blood draw to preserve intracellular signaling states for analysis by mass cytometry. Paired PB samples were collected from patients (n=32) in the NordCML007 trial before and one hour after the first per oral dasatinib dose on day one (t0, t1), and after three months of daily dasatinib, before and one hour after the daily per oral dasatinib (t2, t3). Similarly, leukocytes were collected and preserved from patients (n=49) in the ENEST1st trial before and three hours after the first nilotinib dose (et0, et1) on day one. The different intervals between before and after samples reflected the median time to maximum serum concentration of dasatinib (∼1 hr) and nilotinib (∼3 hrs). The patient samples, together with healthy donor samples (HDPB; n=17, HDBM; n=19), were stained with a panel of 43 metal-conjugated antibodies enabling deep immunophenotyping and profiling of BCR::ABL1 related signaling pathways. In addition, cationic granular proteins of eosinophils and neutrophils were labelled by cisplatin (^Pt^195) through charge-based interactions (49) and used in immunophenotyping alongside antibody derived data (**Supplemental Experimental Procedures**). We acquired high-dimensional data from a total of 44 million single cells from patient and healthy donor samples. Firstly, the main immune populations were identified using Cytosplore (HSNE(50)) clustering and secondly, a selection of these populations were re-clustered using PARC(51) and a tailored selection of markers to further increase phenotypic resolution (**Figure–1B,C**). After manual curation and annotation of the clusters, 37 unique immune populations were identified. The data from each clinical trial were analyzed independently but following the same bioinformatic pipeline, and no batch correction was performed (**Figure–1D**). Differential testing using proportions of immune populations (%, compositional data) may be biased, as a change in the absolute counts of one cell type (or cluster) will affect the proportions of all clusters.(52) To avoid bias from dasatinib-induced acute but transient lymphocytosis (27, 28) affecting cluster proportions, we used routine white blood cell (WBC) absolute counts (10^9^ cells/L) obtained at study visits to estimate absolute counts from mass cytometry cluster proportions (%). The mass cytometry-derived absolute cluster counts (hereafter eCounts) and routine WBC absolute counts were closely correlated in both clinical trials (**Supplemental figure–S1**). We next used log₂-transformed ratios of eCounts to assess immune cell mobilization following per oral dasatinib and nilotinib (**Figure–1E**). One hour after dasatinib dosing, the most pronounced mobilization was observed in B cells and subsets of NK cells and monocytes, particularly after 3 months of treatment, with cytotoxic T cell subsets showing more modest changes. A similar pattern was observed after the first dasatinib dose on day one, although the log₂-ratio of mobilization was generally lower and did not include monocyte subsets. In contrast, nilotinib treatment did not induce mobilization of any major immune cell populations as has been previously observed.(27) The only exception was a modest but statistically significant reduction in non-classical monocytes three hours after the first dose.

**Figure 1.**
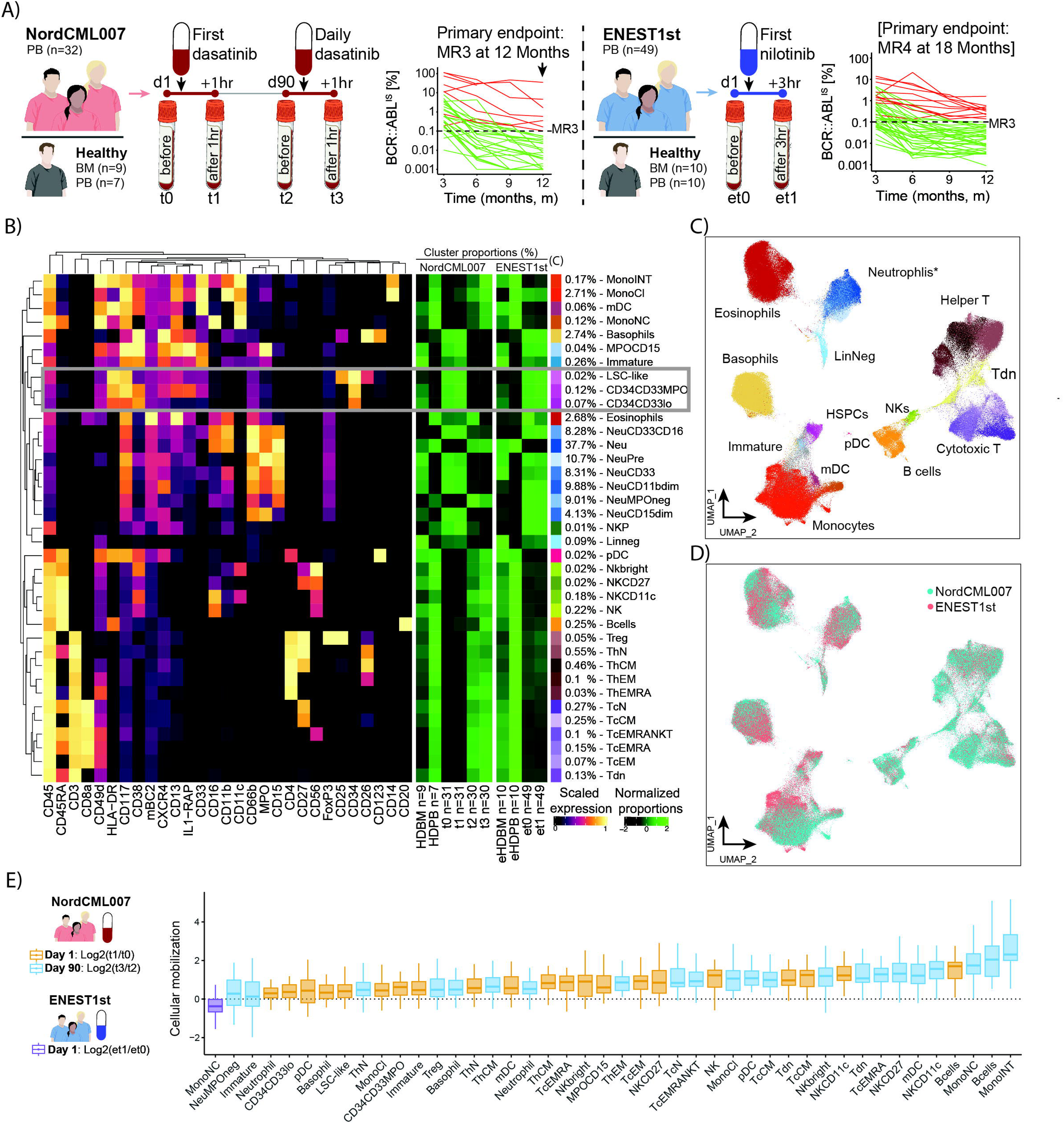
Experimental design and leukocyte mobilization. (A) Experimental design of longitudinal peripheral blood (PB) sampling from, and BCR::ABL1^IS^ monitoring during the first year of treatment in *de novo* patients enrolled in the clinical trials NordCML007 and ENEST1st. Healthy donor PB (HDPB) and bone marrow (HDBM) samples were analyzed alongside each cohort of longitudinal patient samples. (B) Heatmap (left) showing the normalized median expression of proteins used for deep immunophenotyping in 37 immune populations identified by HSNE and PARC clustering. The right green heatmaps show normalized proportions (%) of each cluster in longitudinal patient and healthy donor samples. (C) Uniform manifold approximation and projection (UMAP) representation of single cells from both patients and healthy donors colored by immune population as in B. *The neutrophil clusters were sub-sampled to 10% prior to UMAP-analysis to improve visualization. (D) UMAP as in (C), but colored by clinical trial dataset (NordCML007, ENEST1st). (E) Log_2_ fold change in eCounts of immune populations between t0 and t1, t2 and t3 (NordCML007), and et0 and et1 (ENEST1st), for each patient. Only populations with a significant FDR-adjusted *p*-value (paired *t*-test) are shown.

### Longitudinal deep immune and signaling profiles of CML compared to healthy donors

We next wanted to characterize the dysregulation of PB cells from CML patients before the start of treatment. We therefore compared the composition of immune populations (eCounts), the BCR::ABL1 related signaling pathways and immune-activation state (CD38 and HLA-DR expression) with HDPB.

Since the myeloid cells are expected to express the BCR::ABL1 fusion-protein dringin the disease, this cell compartment was analyzed firstWe found, as expected, that samples from untreated CML patients in both trials (t0 and et0) exhibited significantly increased eCounts of most myeloid cell populations, including basophils, eosinophils, neutrophil subsets, classical monocytes, and dendritic cells compared to PB samples from healthy donor controls (**Figure–2A and B**). The level of pSTAT3 Y705 in the neutrophil subsets was significantly reduced, indicating that these cells, even in seemingly mature and CD16 expressing neutrophils, were less mature than in the phenotypically similar HDPB counterparts (**Figure–2C,D**).(53) Notably, the global level of phosphorylated tyrosine residues (clone: p-Tyr-100) was significantly increased in the basophils. Following three months of dasatinib treatment in the NordCML007 trial, both signaling state and eCounts in most myeloid populations were normalized and comparable to HDPB. There were, however, some exceptions with a significantly reduced eCount of neutrophils, basophils, monocytes, and dendritic cells, which is consistent with myelosuppression, a well-known side effect of TKI treatment.(54, 55) Neutropenia, both grade 2 and 3-4, was detected in 20% of patients in the NordCML007 trial the first 12 months of treatment(56).

**Figure 2.**
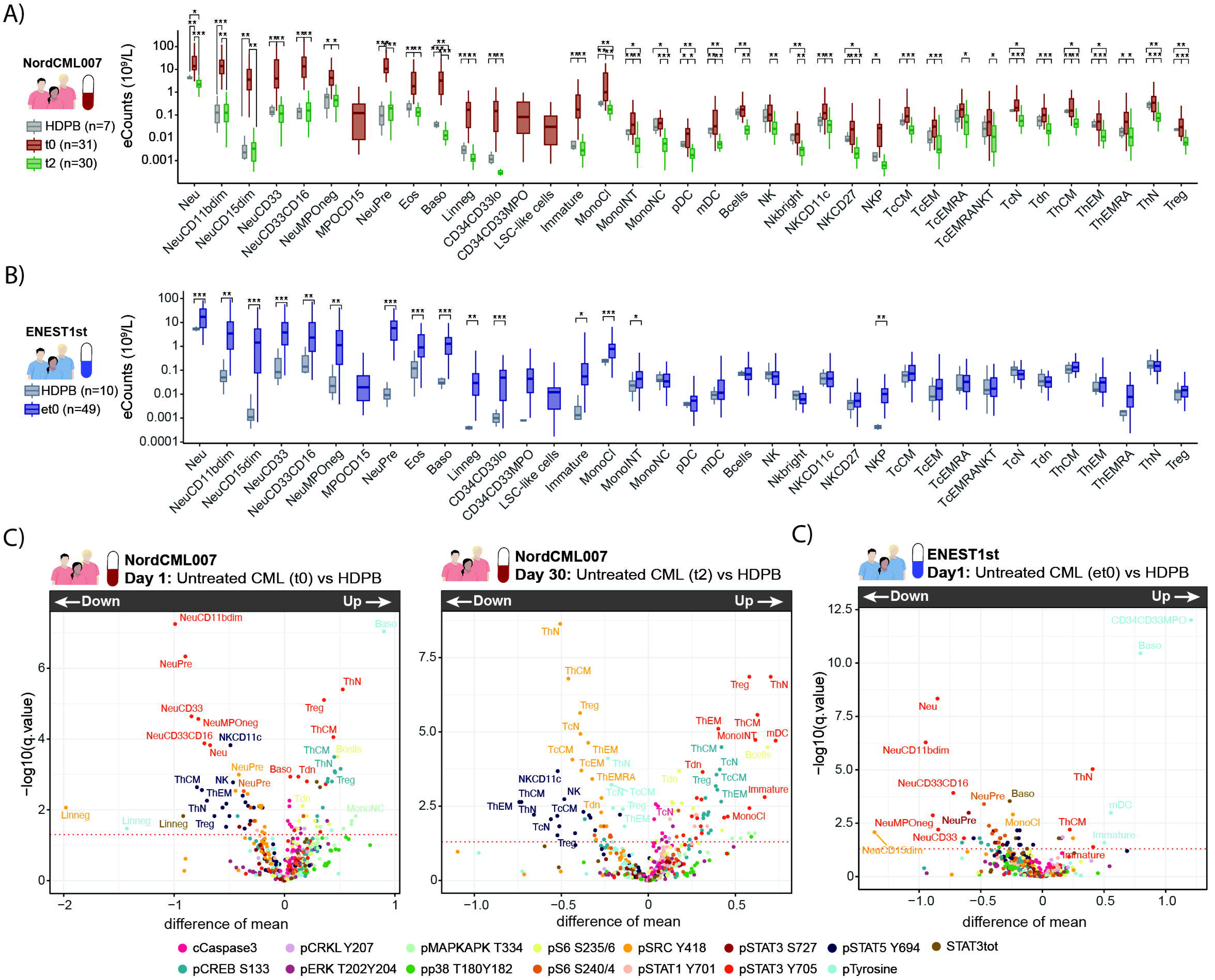
Longitudinal deep immune and signaling profiles of CML. (A) Boxplots showing eCounts of immune populations from patient samples before treatment: dasatinib day 1 (red) and day 90 (green) from NordCML007 trial. Healthy donor peripheral blood (HDPB) samples shown in grey. Statistical comparisons used false discovery rate (FDR) adjusted grouped two-sided t-test (patient vs. HD samples) and paired two-sided t-test (day 1 vs. day 90). (B) Same as in (A) but showing eCounts of immune populations from patients in the ENEST1st trial, before first nilotinib treatment on day 1(et0, blue). Healthy donor peripheral blood (eHDPB) samples shown in grey. (C) Average difference in intracellular signaling states between HDPB and patient samples for each cell type, with FDR adjusted grouped two-sided t-test (patient vs. HDPB) on the y-axis. Differences are shown for samples collected before dasatinib treatment on day 1 (t0, left), or on day 90 (t2, right), from patients in the NordCML007 trial. (D) Same as in (C) but differences are shown for samples collected before dasatinib treatment on day 90 (t2) from patients in the NordCML007 trial, i.e patient vs HDPB All comparisons are based on grouped *t*-tests with for multiple testing. Significance levels are indicated as follows: *Padj* ≤ 0.05: *, *Padj* ≤ 0.01: **, *Padj* ≤ 0.001: ***, *Padj* ≤ 0.0001: ****.

Although the CML-related dysregulation in the myeloid populations in both cohorts was similar before starting treatment, we found that the lymphocyte compartment was more dissimilar. In untreated samples from patients enrolled in the NordCML007 trial, we observed an increased eCount of several lymphocyte subsets, including CD8^+^ central memory (CM) T cells and regulatory T cells (Tregs). Many T cell subsets also exhibited significantly elevated levels of both pCREB S133 and pSTAT3 Y705, and a lower level of pSTAT5 Y694 was observed in both T and NK cells. Overall, we did not make the same observations in the patients in the ENEST1st trial, except in an increased eCount of NK precursors (CD56^-^) and a significantly elevated level of pSTAT3 Y705 in Naïve and CM T helper cells. The eCounts from the HDPB samples collected alongside the NordCML007 (n=7) and ENEST1st (n=10) patient samples were highly consistent, with only a small but significant difference in mDCs (**Supplemental figure–S2**).

Similarly to the myeloid cell subsets after three months of daily dasatinib in the NordCML007 trial, the eCounts of multiple lymphocyte subsets were significantly reduced after dasatinib treatment, most notably B cells, CD56^++^ NKs, CD4^+^ Naïve T cells, and Tregs. Furthermore, a sustained elevation in pCREB S133 and pSTAT3 Y705 levels, along with a reduced pSTAT5 Y694 in T and NK cells was observed, in addition to a decrease in pSRC Y418. The neutrophils showed a clear increase in pSTAT3 Y705, but not pSTAT3 S727, after three months in the various neutrophil populations to the level of healthy donor neutrophils.

Finally, we investigated the level of CD38 and HLA-DR on T and NK cells as a measure of their activation states (**Supplemental figure–S3**). In both patient cohorts, a significantly increased CD38, but not HLA-DR, expression in T helper effector memory (ThEM) cells and reduced expression on Naïve T helper cells were found. The CD38 expression on Naïve T helper cells was further reduced after three months of treatment with dasatinib, while the increased level on ThEM was normalized.

### CML Stem like cells are characterized by increased levels of phosphorylated tyrosine residues including pSTAT5 Y694

Leukemic stem and progenitor cells are of particular interest as their quiescence and resistance to TKI therapy are likely the source of suboptimal response during treatment and relapse after TKI cessation.(40, 41) In the UMAP embedding of cells from patients in both cohorts and healthy donors (**Figure–1C**), we noticed two islands of cells expressing CD33 close to the monocyte subsets, where the most distant cells all expressed CD34. These intermediate and CD34 expressing cells were subjected to a new UMAP analysis to evaluate their immunophenotypic composition in detail (**Figure–3A**). This UMAP analysis clearly showed two distinct islands of cells, separated by CD34 expression. The CML leukemic stem cell markers CD25 and CD26 were co-expressed on a subset of CD34-expressing cells with lower levels of CD38, and these cells were highly enriched in the LSC-like population (CD34^+^CD38^low^CD25^+^). These cells were 47-fold more frequent in leukemic samples compared to healthy donor samples. The proportions of the three CD34^+^ populations were comparable between the two cohorts of patients **(Figure–3B, C)**.

**Figure 3.**
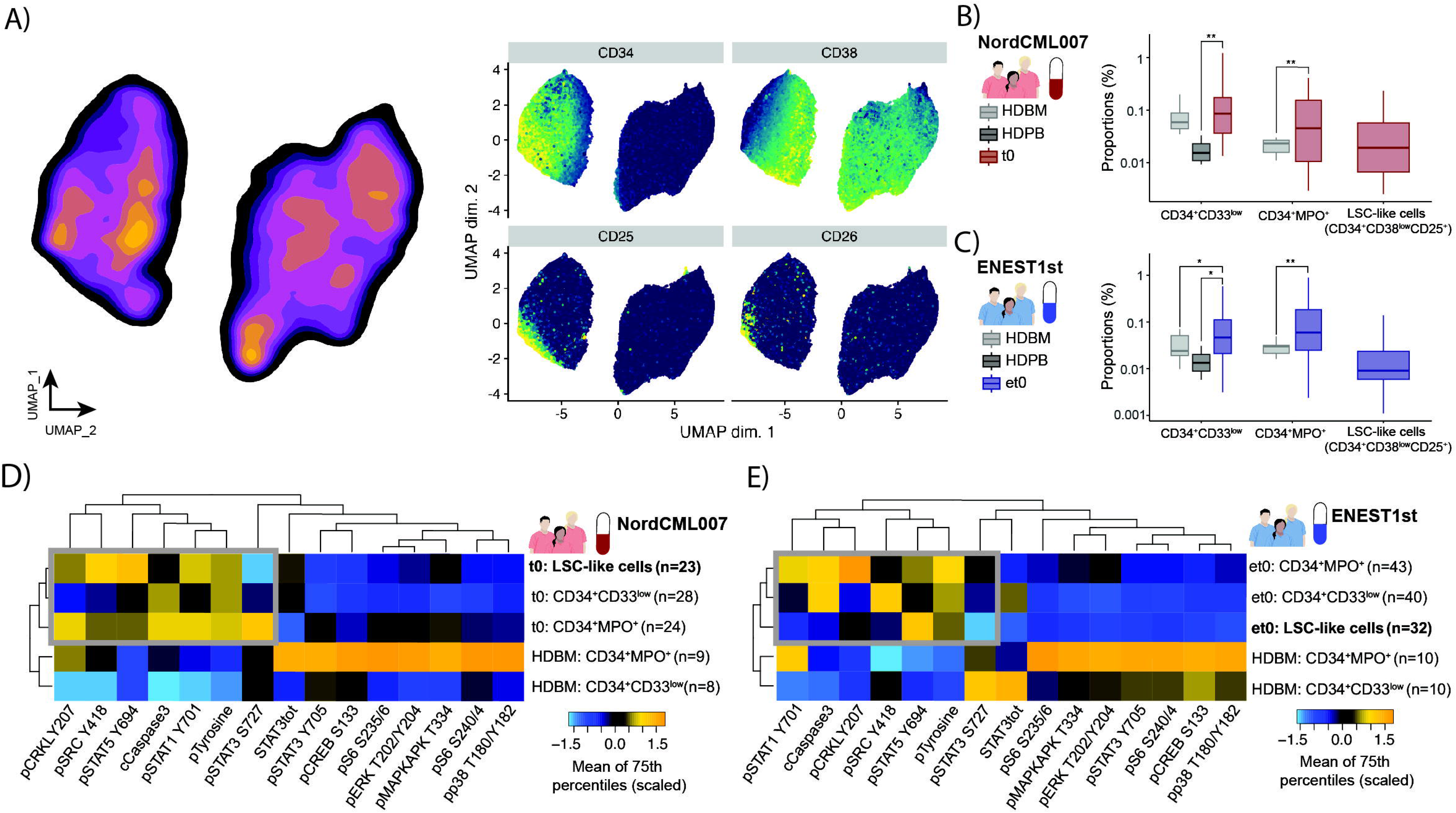
CML stem-like cells display elevated levels of phosphorylated tyrosine residues and pSTAT5 Y694. (A) UMAP embedding of isolated CD33⁺ intermediate and CD34⁺ cells (left). Single-cell expression of selected surface markers with each marker scaled independently (right). (B) Proportions (%) of CD34⁺ cell clusters in samples collected before dasatinib treatment on day 1 (red) and day 90 (green) from patients enrolled in the NordCML007 trial. (C) Proportions (%) of CD34⁺ cell clusters in samples collected before nilotinib treatment on day 1 from patients in the ENEST1st trial. In both B and C, healthy donor peripheral blood (HDPB) and bone marrow (HDBM) samples are shown in shades of grey for comparison. (D) Heatmap showing the scaled average 75th percentile expression of intracellular signaling markers associated with BCR::ABL1 signaling across the CD34⁺ cell clusters. All comparisons are based on groped *t*-tests with false discovery rate (FDR) correction for multiple testing. Significance levels are indicated as follows: *Padj* ≤ 0.05: *, *Padj* ≤ 0.01: **, *Padj* ≤ 0.001: ***, *Padj* ≤ 0.0001: ****.

Next, we characterized the LSC-like population by comparing the state of intracellular signaling in all CD34^+^ populations. For each of the intracellular markers and each CD34^+^ population, we calculated the average 75^th^ percentile expression for healthy donor BM samples and the untreated CML samples (**Figure–3D, E**). We found that the signal transduction profiles of CD34^+^ populations from untreated CML patients were clearly unique compared to phenotypically similar cells from HDBM, and consistent between patient samples from the two trials. As expected in cells expressing BCR::ABL1, the CD34^+^ patient populations all had a higher level of phosphorylated tyrosine residues compared to the healthy clusters. The CD25 and CD26 expressing LSC-like cells were further distinct by a higher level of pSTAT5 Y694 in addition to a lower level of pSTAT3 S727. Interestingly, cleaved caspase 3 was also lowest in in the LSC-like cells.

### Signaling kinetics of TKIs after first per oral dosing

Having confirmed that our early longitudinal sampling captured dasatinib-induced changes after one hour, exemplified by acute immune cell mobilization (**Figure–1E**), we next investigated intracellular signaling changes induced by TKI treatment. We began by examining global levels of phosphorylated tyrosine residues in basophils and CD34^+^ populations (**Figure–4A,B**). Compared to HDPB samples, relative phosphorylated tyrosine levels were 3.26±0.31 and 5.68±1.14-fold higher in basophils from untreated CML patients, for nilotinib and dasatinib, respectively, consistent with constitutive BCR::ABL1 activity driving elevated tyrosine phosphorylation. Strikingly, in both patient cohorts, we observed a highly significant decrease in phosphorylated tyrosine one hour after the first dasatinib (p_adj_ = 2.42×10^-11^) and three hours after the first nilotinib tablet (p_adj_ = 8.64×10^-11^). Interestingly, in the basophils after 3 months of daily dasatinib treatment, dasatinib also induced a significant (p_adj_ = 1.32×10^-4^) reduction of phosphorylated tyrosine residues.

**Figure 4.**
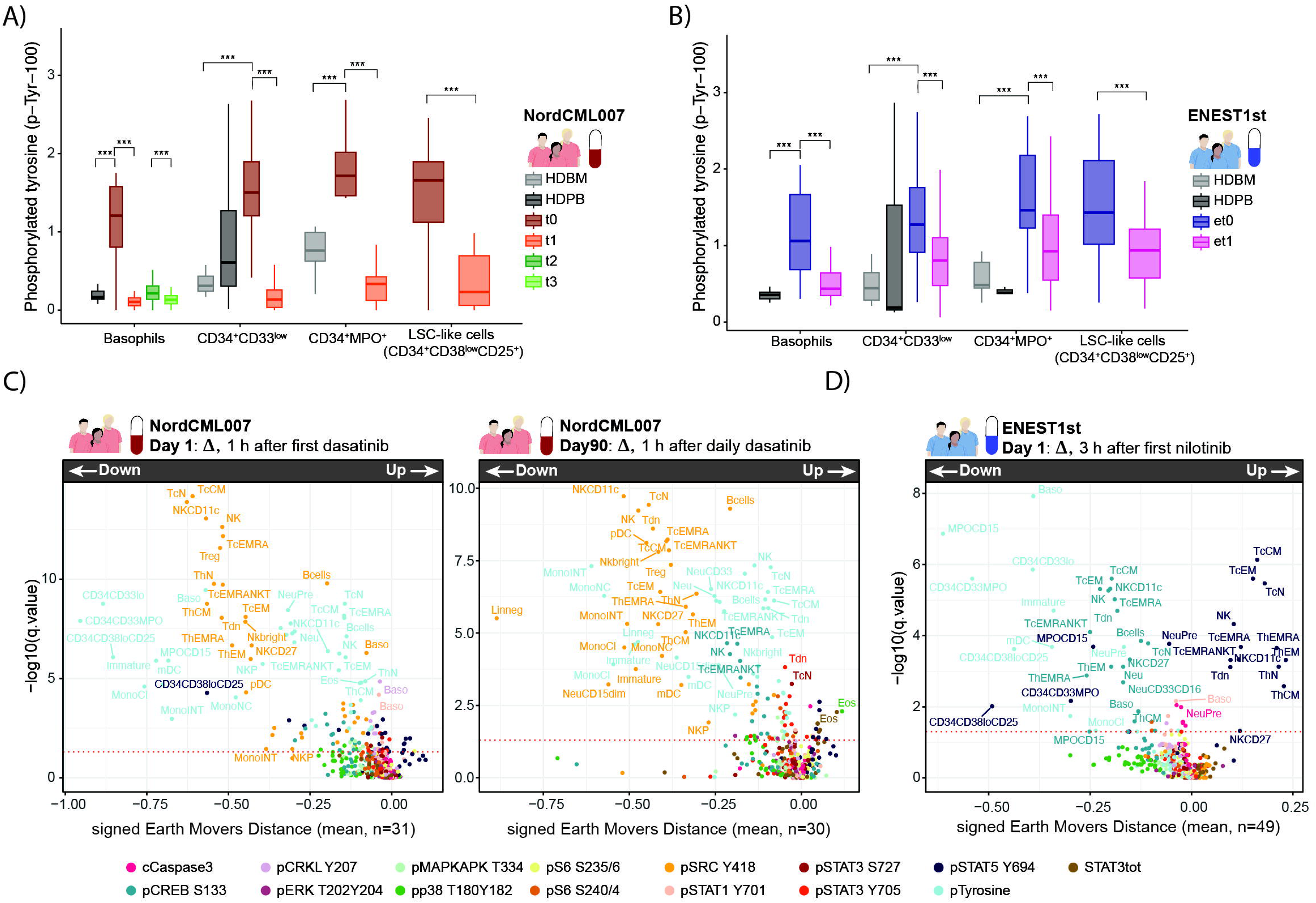
TKIs inhibits signaling hours after first per oral dosing. (A-B) Boxplots show the 75th percentile measurement of phosphorylated tyrosine residues using the P-Tyr-100 antibody clone in basophils and CD34⁺ cell clusters from patients with CML across longitudinal patient samples from patients treated with dasatinib and nilotinib, respectively. The 75th percentile of phosphorylated tyrosine residues from same populations from HD healthy donor peripheral blood (HDPB) and bone marrow (HDBM) samples are shown in shades of grey. (C-D) Signed Earth Movers Distance (sEMD) measures of changes in signalling one (dasatinib; day 1; left and day 90; right) and three hours (nilotinib) after first dose of tyrosine kinase inhibitor. These sEMD measures of changes of signalling have been visualized with an FDR-adjusted and paired t-test analysis using 75th percentiles on the y-axis. All comparisons are based on paired t-tests with false discovery rate (FDR) correction for multiple testing. Significance levels are indicated as follows: padj ≤ 0.05: *, padj ≤ 0.01: **, Padj ≤ 0.001: ***, padj ≤ 0.0001: ****.

The level of tyrosine phosphorylation was higher in the untreated patient CD34^+^ populations compared to both the basophils in the patient samples and compared to CD34^+^ populations from healthy donors, providing indirect evidence of BCR::ABL1 expression.(57, 58) Additionally, we also observed a highly significant reduction in phosphorylated tyrosine one hour after the first dasatinib and three hours after the first nilotinib tablet in the CD34^+^ populations, including the LSC-like cells (dasatinib: p_adj_ = 1.37×10^-7^, nilotinib: p_adj_ = 1.22×10^-5^).

To characterize the broader effect of the TKIs on intracellular signaling associated with BCR::ABL1 we used signed earth movers distance measure (sEMD) combined with an FDR adjusted and paired two-sided t-test of 75^th^ percentile measurements in **Figure–4C,D**. Similar to basophils and LSC-like cells, we observed a broad reduction in phosphorylated tyrosine levels across most immune populations, with the most pronounced effects in myeloid lineage cells (monocytes and other CD34^+^ populations) compared to lymphoid cells. A hallmark of BCR::ABL1-inhibition is the downregulation of pSTAT5 Y694 in CML stem cells and cell lines.(59, 60) In our patient samples, hours following per oral TKI dosing we observed a significant inhibition of pSTAT5 Y694 in all three CD34^+^ cell populations in patients treated with both dasatinib and nilotinib, with the most prominent downregulation observed in the LSC-like cells. In a population of cells with a phenotype resembling pre-neutrophils (CD66b^+^CD34^-^CD117^+^CD38^+^CD49d^+^) we observed a significant reduction in pSTAT5 Y694 in both dasatinib and Nilotinib treated patients.(61-63) Dasatinib treatment also resulted in a modest but significant downregulation of pCRKL Y207, pSTAT1 Y701 and pSRC Y418 in basophils, similar previous observations 3 hours after first nilotinib(46), and in most neutrophils populations (**Supplemental figure–S4**).

In the lymphoid compartment, we measured a downregulation of pSRC Y418 following dasatinib treatment in most cell subpopulations, but most strikingly in T and NK cell subsets. This is likely reflecting dasatinib inhibition of SRC-family kinases and not ABL1, and no regulation of pSRC Y418 was observed in nilotinib treated patients. Interestingly, inhibition of pSRC Y418 was also readily measurable one hour after dasatinib dosing in samples collected after three months of treatment. In the nilotinib treated patients (**Figure–4D)** we were able to validate previous results showing an increase in pSTAT5 Y694 alongside a reduction in pCREB S133 in T and NK cells.

### Correlated Immune and Signaling Features Reveal Systemic Effects of TKIs

To investigate the systemic longitudinal effects of TKIs on peripheral blood leukocytes, we constructed networks of immune features using Spearman correlation and UMAP dimensionality reduction. In these networks, each immune feature is represented by a node that is defined by the data type (baseline or longitudinal change) of a specific variable (eCount, CD38 expression, BCR::ABL1-related signaling) in an immune population. Only significantly correlated immune features are plotted and these are connected by p_adj_ weighted edges. Nodes with similar patient profiles are proximal to each other on the UMAP.

We first built a network from the ENEST1st patient data (**Figure–5A**) that included measurements from all immune populations at both baseline (BL, et0, red) and change after three hours of nilotinib (Ll, et1–et0, yellow) on day 1. The UMAP organization was primarily driven by data type, with strong correlations observed within baseline and change after three hours of nilotinib. However, there were some correlations between baseline and change after three hours of nilotinib immune features. Immune cell populations generated additional UMAP structure, with neutrophil subsets mapping together and separately from heterogenous communities of lymphocytes. Analysis of signaling markers revealed highly organized but more local patterns, where heterogeneous lymphocyte subgroups exhibited similar baseline or treatment-induced changes in signaling profiles. This was especially evident when examining the correlation between pMAPKAPK T334, pp38 T180/Y182 and pCREB S133.

**Figure 5.**
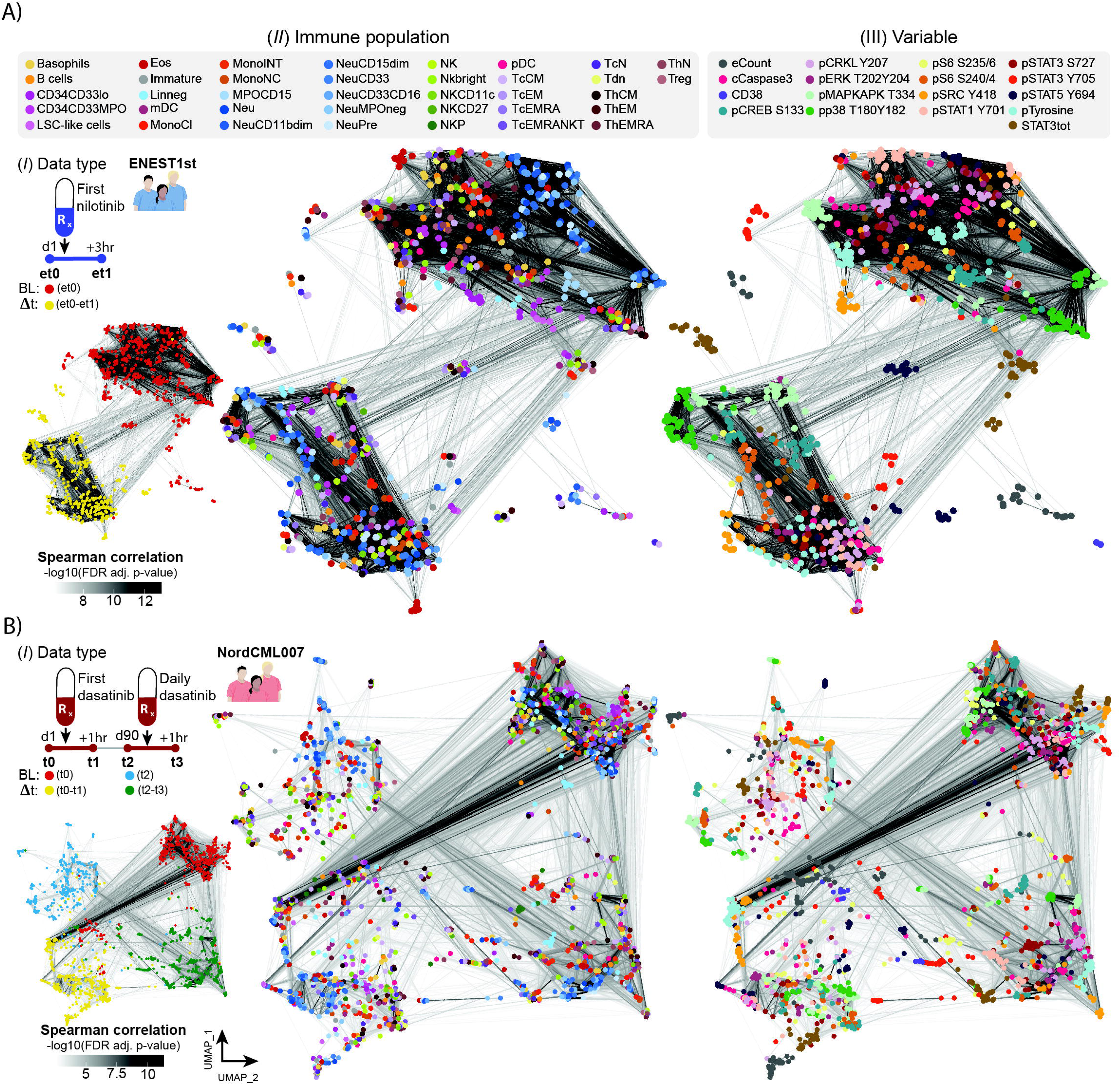
Correlated Immune and Signaling Features Reveal Systemic Effects of TKIs. (A-B) Network visualization of correlated immune features across nilotinib (*above*) and dasatinib (*below*) treatment patients. Each node represents an immune feature positioned using UMAP dimensionality reduction, colored by data type (*I*, baseline or longitudinal change), immune population (*II) and* variable (*III,* eCount, CD38 expression, BCR::ABL1-related signaling). Edges connect significantly correlated features (FDR-adjusted Spearman p-value < 0.05), with edge transparency reflecting correlation strength. Nodes with similar profiles across patients cluster together

We then analyzed the NordCML007 trial patient data (**Figure–5B**), capturing immune features across multiple longitudinal samples: baseline on day 1 (BL, t0, red) and day 90 (BL, t2, blue) and treatment-induced changes after one hour of dasatinib on day 1 and day 90 (Δt1–t0, yellow, and Δt3–t2, green). As we observed in the ENEST1st network, the UMAP was broadly organized by data type, and there was considerable correlation between nodes both within and across data type at the start of TKI treatment, and after three months. The most prominent feature of the network was a highly significant and positive correlation between baseline (t0) and change one hour after dasatinib on day 1 (Δt1–t0) of phosphorylated tyrosine residues across various immune populations. This correlation pattern was absent in the ENEST1st network, likely due to differences in dosing regimens between the two studies. In the ENEST1st trial, samples were collected before and 3 hours after a single 300 mg dose of nilotinib (half of the standard 300 mg twice-daily regimen), possibly resulting in incomplete BCR::ABL1 inhibition. In contrast, NordCML007 patients received the full 100 mg once-daily dose of dasatinib. The incomplete target inhibition in the nilotinib cohort may explain the absence of the strong baseline-to-change correlations observed with full-dose dasatinib, suggesting that the magnitude of kinase inhibition influences the coordinated signaling response across immune cell populations. Overall, our feature-network highlighted that groups of different cell types were closely located in the UMAP based on similar baseline levels or changes in intracellular protein phosphorylation. These local structures of cell types then formed an integrated network of highly correlated measurements between nodes in signal transduction pathways, like pp38 T180/Y182 and pMAPKAPK T334. From our networks we can see a clearly shared and coordinated immune response to TKI treatment that may be dose dependent in phenotypic diverse cell subsets but broadly delineated by BCR::ABL1 positive and negative cell types.

### Responders to dasatinib and nilotinib are predicted by unique sets of immune features

We next asked whether our immune features were predictive of treatment response as measured by *BCR::ABL1* mRNA levels (BCR::ABL1^IS^) every three months during the first year of treatment. To enable direct comparison between the dasatinib and nilotinib cohorts, we standardized our analysis to immune features derived from equivalent early timepoints: baseline measurements before TKI initiation (t0 for dasatinib, et0 for nilotinib) and acute treatment-induced changes measured one or three hours after the first dose (Δ, t1–t0; Δ, et1–et).

To get an overview of the correlated immune features, we first performed Spearman correlation analysis of immune features and BCR::ABL^IS^ measurements the first year of treatment in each cohort. Overall, the number and strength of significantly correlated immune features in nilotinib-treated patients were lower than in the dasatinib-treated patients (**Figure–6A, B**). Features related to immune activation (pERK ½ in ThEM cells) and high leukemic burden (eCounts of myeloid and CD34^+^ cells and pSTAT3 Y705 in neutrophils) was the dominant immune features correlating with response to dasatinib. In contrast, in nilotinib-treated patients immune features correlating response were largely driven by the LSC-like cell population. Specifically, baseline levels of cleaved Caspase-3, pCRKL Y207 and STAT3 (total and phosphorylated) in LSC-like cells before starting treatment with nilotinib were negatively correlated with BCR::ABL^IS^. Whereas the increase in cleaved Caspase-3 LSC-like cells three hours after first nilotinib tablet was positively correlated with BCR::ABL1^IS^. In addition to the LSC-like population, the change in eCounts of NK progenitors (NKP) after three hours and baseline classical monocytes (MonoCl) we all positively correlated with BCR::ABL1^IS^.

**Figure 6.**
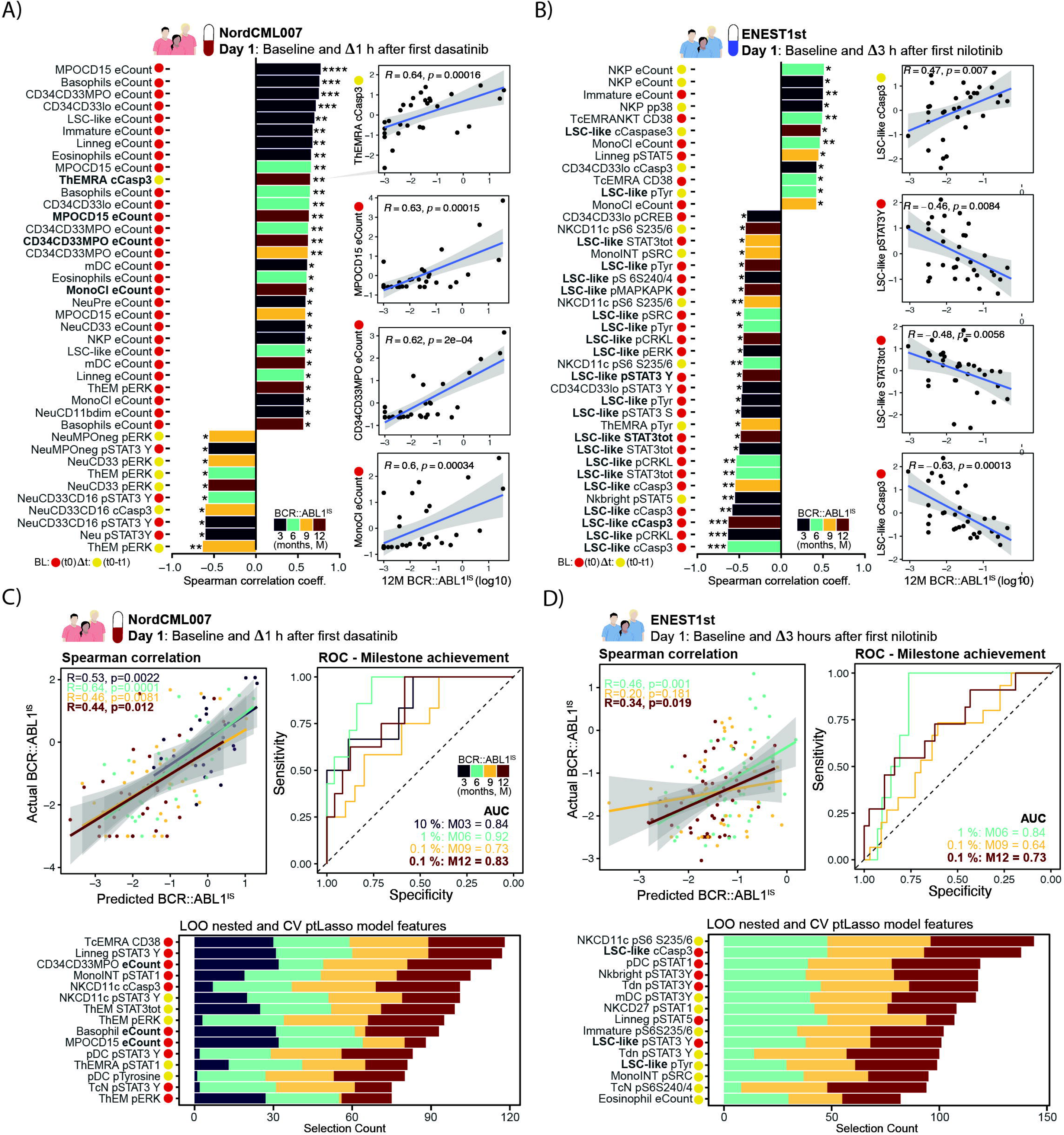
Responders to dasatinib and nilotinib are defined by unique sets of immune features. (A-B) Top 30 Spearman correlations (ranked by correlation coefficient R) between immune features and BCR::ABL1^IS^ [%] levels at 3-, 6-, 9-, and 12-months post-treatment initiation in dasatinib-treated patients. Immune features extracted from baseline samples (red, t0/et0) and changes from baseline to 1 hour (dasatinib) or 3 hours (nilotinib) after first TKI dose (yellow). (C-D) Predictive modeling of BCR::ABL1^IS^ using pretrained LASSO logistic regression on the same data as above. Model performance evaluated by Spearman correlation between predicted and observed BCR::ABL1^IS^ and Receiver operating characteristics and Area under the Curve (AUC) statistics for classifying responders at milestone timepoints (BCR::ABL1^IS^ ≤10% at 3 months, ≤1% at 6 months, ≤0.1% at 9 and 12 months). Selection frequency of immune features during the double-layered leave-one-out nested and cross-validated pretrained lasso analysis.

To move beyond pairwise associations, we built a predictive model of treatment response using a pretrained LASSO variant (ptLasso). LASSO supports multivariate modeling while selecting a minimal set of informative features, producing interpretable, generalizable models. Pretraining captured relationships common to all BCR::ABL1^IS^ outcomes in the first treatment year, followed by month-specific fine-tuning to enhance individual predictions.(64) Model fitting and performance were evaluated with leave-one-out nested cross-validation to ensure unbiased estimates.(65) Using early immune and signaling features, we trained models within each clinical trial to predict BCR::ABL1^IS^ every three months during the first year of treatment. Model performance, evaluated by Spearman correlation, revealed strong agreement between predicted and observed BCR::ABL1^IS^ values in both NordCML007 (12 months, R = 0.44, p = 1.2 × 10^⁻2^) and ENEST1st (12 months, R = 0.34, *r* = 1.9 × 10⁻^2^). We further assessed classification performance by predicting optimal versus suboptimal responders at each milestone timepoint (BCR::ABL1^IS^ ≤10% at 3 months, ≤1% at 6 months, and ≤0.1% at 9 and 12 months; **Figure–6C,D**). The models achieved high discriminative performance, with AUCs of 0.83 (NordCML007) and 0.73 (ENEST1st) at 12 months. Notably, many of the top-ranked features selected by the model overlapped with those identified in the Spearman analysis, reiterating that the features predicting the response to dasatinib and nilotinib are different.

## Discussion

There is a fundamental temporal disconnection between the biology of targeted cancer therapy and current response evaluation methods. While pharmacodynamic studies demonstrate that cellular responses to targeted therapy can be measured within hours through intracellular signaling monitoring (46, 48, 66, 67), clinical response evaluation in CML relies on assessments performed every three months (BCR::ABL1^IS^).(9) Here, we show that immune features captured within hours of initiating TKI treatment predict long-term therapeutic outcomes at the individual patient level, despite similar group-level efficacy between dasatinib and nilotinib **Supplemental figure–S5**).(68-70) These rapid, single-cell measurements of intracellular signaling states across distinct cellular subsets, together with tumor and immune cell composition, reveal coordinated patterns of TKI response. Our findings open a new avenue in ultra-rapid response evaluation of targeted cancer therapy.

The interactions between leukemic cells and the immune system are critical determinants of treatment outcomes in cancer.(71-73) We find that that immune features measured hours after dasatinib initiation that correlated with treatment response were dominated by leukemic burden, neutrophil maturity (pSTAT3 Y705), and immune cell activity (pERK ½ in T cells and NK cells). These features reflect the broad kinase selectivity of dasatinib beyond ABL1, particularly its targeting of SFK-kinases, and provide additional evidence for the importance of dasatinib’s immunomodulatory mechanisms (in?) cells.(23, 25, 74, 75) Huuhtanen and co-workers analyzed samples from the same NordCML007 trial patients by performing comprehensive immune profiling using flow cytometry, scRNA-seq, and functional assays on cryopreserved PBMCs.(22) While focused primarily on cellular proportions and functional changes following IFN-α addition to dasatinib therapy, baseline CD107^+^GZMB^+^ NK cells correlated with BCR::ABL^IS^ after 12 and 18 months of treatment. This is consistent with a feature in our machine learning model, identifying the change in pSTAT3 Y705 in CD11c^+^ NK cells correlating to response after 12 months of treatment.

In contrast to the effect of dasatinib, the effect of nilotinib was predominantly centered on CD34^+^CD38^low^CD25^+^ and CD26^+^ LSC-like cells and their intrinsic signaling states (cCaspase-3, STAT3 [total and Y705], and global tyrosine phosphorylation changes). The high selectivity of nilotinib to BCR::ABL1, and consequently its minimal impact on the immune system, suggests that nilotinib treatment outcomes are more dependent on the LSCs survival pathways alone.(76, 77) When machine learning approaches were applied to the same data to predict individual patient outcomes during the first year of treatment, there was substantial overlap between the immune features selected by the model and those identified through Spearman correlation analysis. These findings suggest that baseline characteristics and single-cell immune and signaling profiles could be leveraged to identify which patients are more suitable candidates for dasatinib versus nilotinib, thereby enabling personalized CML selection of TKIs. E.g., patients with a high leukemic burden and increased cCaspase3/pCRKL/pSTAT3 signal should start dasatinib, while patients with the opposite profile should start nilotinib. We hypothesize that our prediction of response to TKI can be leveraged to guide selection of the optimal TKI at start of treatment and will increase the proportion of patients eligible to trial discontinuation of treatment. This needs to be tested in clinical trials.

Several factors influence the on-target effects of TKIs in patients, including the actual dose ingested by the patient or other drugs taken concomitantly, and pharmacokinetic factors such as GI absorption and hepatic metabolism. When we compared the reduction in basophil tyrosine phosphorylation levels after the first dose of TKI, we observed differences in patients treated with dasatinib and nilotinib. The residual basophil tyrosine phosphorylation, 1 hour after dasatinib, was below or comparable to HDPB (mean ratio: 0.55±0.12). In the patients treated with nilotinib however, we observed higher residual tyrosine phosphorylation compared to HDPB (mean ratio 1.57±0.17). This could be explained by the fact that nilotinib was given as 300 mg twice daily, and our peripheral blood samples were collected from patients who had only received one of the two daily doses of nilotinib. This observation indicates that we can measure the degree to which a TKI is inhibiting intracellular signaling in cellular subsets.

## Limitations of this study

An important limitation in this study is the relatively small sample size. In ENEST1st clinical trial the collection of samples was planned from 200 patients, but we only obtained usable samples from 50 patients from centers across European centers. In NordCML007 we collected and analyzed samples from 32 out 40 patients enrolled in centers in the Nordic countries. In these studies, we used the BD Lyse/Fix (BD Phosflow; 37°C water bath, centrifugation, storage -80°C) for the preservation of white blood cells. Recently, more simple preservation buffers have become commercially available (Smart tubes, Cytodelics) significantly easing the sample preservation to only 10 mins following blood draw. We expect new sampling tube technology to significantly increase the feasibility of larger studies of immediately preserved full blood.

The peripheral blood samples were collected from patients in two clinical trials with different inclusion criteria for prior treatments allowed before starting study drug. The NordCML007 trial only allowed prior hydroxyurea (HU) for up to 60 days, whilst ENEST1st allowed TKI treatment up to 3 months and HU for up to 6 months. In the ENEST1st trial, only one patient had received prior TKI-treatment and there was no significantly increased duration (days) of HU (**Supplemental figure–S6**) compared to the NordCML007 trial (Wilcoxon rank sum test P = 0.2592).

Systematic technical variation from sample preservation, antibody labelling and instrumentation is a significant hurdle for clinical application of longitudinal studies using high-dimensional single-cell technologies. We partly mitigated this in our sample preparation by staining all samples on the same day using the same frozen and aliquoted antibody master mix and storing the labelled samples at -80°C for data acquisition. The data from the NordCML007 samples were acquired on a Helios mass cytometer within weeks, while ENEST1st samples were collected a year later using a CyTOF XT instrument. Despite this, we observed a minimal batch effect (Figure 1D, **Supplemental figure–S2**) and chose to analyze the data without any normalization. This work lays the ground for future clinical implementations where a single longitudinally collected sample can be analyzed independently to generate clinically meaningful data for response prediction—without requiring normalization to large reference cohorts or standardization to a single instrument platform.

## Concluding remarks

We find that dasatinib and nilotinib induces changes in intracellular signaling hours after the first –per-oral dose, allowing for a rapid therapy optimalization. Methodologically, it is feasible to measure the effects of targeted treatment by modifications of protein kinases within minutes or hours after the drug exposure. (78, 79) We suggest that this should be performed with single cell resolution to secure information from TKIs with different profiles of target inhibition. Single cell immune and single cell profiling may in this way be developed into a next generation white blood cell count, providing a precision in diagnostics that matches the precision of targeted therapy.

## Experimental Procedures

### Patients and healthy donor samples

Longitudinal peripheral blood (PB) samples were collected from patients with newly diagnosed chronic phase CML enrolled in the NordCML007 (NCT01725204)(56, 80), and ENEST1st (NCT01061177)(81) clinical trials (patient characteristics: **Supplemental Table S1**). From patients (n=32) in the NordML007 trial, PB samples (N=122) were collected from each patient before and 1 hour after the first dasatinib dose (t0, t1), and before and 1 hour after dasatinib dose after 3 months of daily dasatinib treatment (t2, t3). The NordCML007 trial studied the efficacy and safety of combining dasatinib (100 mg once daily) with pegylated interferon-α2b (PegIFNα), after three months of dasatinib treatment. The PegIFNα was given as 15 µg/week for three months then 25 µg/week for nine months. Our longitudinal sample collection was performed before the addition of PegIFNα. From patients (n=49) in the ENEST1st trial treated with nilotinib (300 mg twice daily), PB samples (N=98) were collected from each patient before and 3 hours after the first nilotinib dose (et0, et1). Although NordCML007 and ENEST1st both enrolled newly diagnosed chronic phase CML, the inclusion and exclusion criterions were different. While the NordCML007 trial did not allow any prior treatment TKIs, and only up to 3 months of prior hydroxyurea (HU) treatment, the ENEST1st trial allowed TKI treatment for up to 3 months prior, and HU for up to 6 months prior. PB (n=17) and bone marrow (BM, n=19) samples from healthy donors were collected after written informed consent (University of Bergen, Norway, ethical committee approval 2012/2247. All samples were preserved for mass cytometry analysis using Phosflow Lyse/Fix buffer (BD, cat# 558049) and immediately frozen in saline for long-term storage at −80°C, as previously described.(46)

#### Barcoding

The same barcoding strategy was applied to patient samples from both trials. Up to 40 samples, including CML, HD, and technical replicates, were barcoded and pooled. To achieve this, two pools of 20 samples each were barcoded and pooled using in-house formulated 20-plex palladium reagents as described by Zunder et al., 2015.(82) These two 20-plex barcode pools were further barcoded and pooled using two unique combinations of three cisplatin isotopes (^194^Pt+^195^Pt and ^198^Pt+^195^Pt), following an approach described in McCarthy et al., 2017.(83) We designed the Pt barcoding such that all cells in all pools were stained with ^195^Pt. Eosinophils take up distinctly more ^195^Pt than other cell types, and was therefore used alongside antibody-derived cell surface marker expression during clustering and dimensionality reduction (**Supplemental Experimental Procedures**).

#### Single cell staining

The two master mixes of antibodies targeting cell surface markers and intracellular protein epitopes (**Supplementary Table S2**) were prepared prior to single-cell staining and stored in aliquots at -80°C.(84) All barcoded sample pools from both trials were stained on the same day using the frozen antibody master mixes, according to a modified version of Standard BioTools’ protocol (Maxpar Cell Surface Staining with Fresh Fix Protocol, 400276 Rev 07).

Briefly, after metal barcoding, cells were thawed, washed and incubated with heparin sodium (100 IU/mL, Wockhardt, Cat # FP1083) and FcR Blocking Reagent (1:5 dilution, Miltenyi Biotec, Cat # 130-059-901) for 20 minutes at room temperature to inhibit unspecific charge-based and FcR-mediated antibody binding.(49) The cells were then stained with the panel of antibodies for cell surface markers (30 minutes, RT), washed in cell staining buffer (CSB, SBT cat # 201068), and permeabilized with methanol (-20°C, Cat #) on ice for 15 min. Further washing in CSB was followed by a second round of blocking with heparin and FcR blocking reagent. Cells were then washed in CSB and stained with the panel of antibodies for intracellular targets (30 minutes, RT). After washing, the cells were resuspended in PBS (SBT, Cat # 201058) + 4% PFA (Thermo fisher, cat # 28906) with a 1:3000 dilution of iridium-intercalator (125 uM, SBT, cat # 201192A) and incubated overnight at 4°C. The next day, cells were washed in CSB and frozen in FBS (?) supplemented with 10% DMSO (?) at -80°C until data acquisition.(85) A pool of single-stained compensation beads was also prepared according to previously published protocols.(86)

#### Data acquisition

The stained and frozen barcode pools from the NordCML007 trial were thawed immediately prior to data acquisition, washed in CSB and Cell Acquisition buffer (CAS, STB, cat # 201240) and re-suspended to a final concentration of approximately 5×10^5^ cells/mL in CAS containing Four element EQ Normalization beads (1:10 dilution, STB Cat # 201078). The data was acquired on a Helios mass cytometer using a wide bore injector at an event rate of **∼** 300 events/second. A Super Sampler (Victorian airships inc.) was used to inject cells to the mass cytometer. The single cell sample suspensions were kept on ice during acquisition.

The stained and frozen barcode pools from the ENESt1st trial were thawed immediately prior to data acquisition, washed in CSB and Cell Acquisition buffer+ (CAS+, STB, Cat # 201244) and left pelleted in flow tubes. The flow tubes were placed in the sample carousel of the CyTOF XT (SBT) and cell pellets were re-suspended to a final concentration of approximately 5×10^5^ cells/mL in CAS+ containing Six element EQ Normalization beads (1:10 dilution, SBT, Cat # 201245) during data acquisition. The data was acquired at an event rate of **∼** 300 events/second.

### Data analysis

#### Pre-processing of single-cell data

The data was normalized using the Fluidigm normalizer software based on the four and six element normalization beads in single cell data collected from samples from the NordCML007 and ENEST1st patients, respectively. The normalized data was cleaned up by manual gating using the Gaussian parameters, Event length and DNA/Ir as outlined in Bagwell et al., 2020.(87) The panel editor tool from the R package *Premessa* was used to harmonize the panels and to delete the BCKG-190 channel. The data was de-barcoded and compensated using the R package *CATALYST*. Data from acquired single stained compensation beads was used for compensation.(86) Final removal of doublets was performed by tailored gating on CD45 vs DNA/Ir, with DNA/Ir channel set to linear. The single cell data was not batch effect corrected.

#### Clustering and dimensionality reduction

Both datasets were clustered and manually annotated using the same overall two-step strategy. The first step, the main cell types were identified using Cytosplore.(50) Then, T cells, Myeloid and dendritic cells, NK cells, hematopoietic stem and progenitor cells (HSPCs) and neutrophils were re-clustered individually using PARC(51) and a tailored selection of cell surface markers (**Supplementary Table S3**). The same nomenclature of cell type labels was used in the annotation of clusters to enable direct comparison between datasets. Uniform Manifold Approximation and Projection for Dimensional Reduction (UMAP, arXiv:1802.034269) was used for dimensionality reduction analysis for visualization of single-cell level data.

#### Estimation of absolute counts from proportions

To estimate the absolute counts of different immune populations identified by mass cytometry, we multiplied the proportion of each population by the white blood cell count (WBC, 10^9^/L) measured on the day of PB sampling. These estimated absolute counts were then adjusted to account for the rapid lymphocyte mobilization increasing the white blood count.(27) For details, see supplemental materials.

#### Comparison of signaling in healthy donors and patient samples

We performed a group-wise comparison between healthy donors and patients before, or after three months of treatment by calculating the difference (Δ) in mean level of each of the BCR::ABL1 related signaling pathways in each immune population for healthy donor samples and in CML patient samples. The level of BCR::ABL1 related signaling was measured in the immune populations as the 75^th^ percentile of the hyperbolic arcsin-transformed dual counts.

#### Change in signaling during treatment

For each patient, to quantify the distance between cell distributions, the Earth Mover’s Distance (EMD) was employed. This metric captures the minimal shift in probability mass required to align one distribution of cellular features with another, reflecting differences in their overall statistical structure. To assess the directionality of these shifts, a signed Earth Mover’s Distance (sEMD) was defined by incorporating the sign of the median difference between perturbed and baseline variants.

### Statistical analysis

We used the t_test() function from the R package *rstatix* to do simple two-sided Student’s t-tests. To perform Spearman correlations, we either used the rcorr() function from the R package *Hmisc* (correlation networks), or stat_cor() from the R package *ggpubr* for simple plotting. Results from all statistical tests, including Spearman correlations, were false discovery rate adjusted using the Benjamini & Hochberg method in the p.adjust() function in base R (p_adj_). We used the cv.ptLasso() function from the R package *ptLasso* to predict response to treatment from mass cytometry data. Receiver Operator Curves and Area Under the Curve (AUC) measurements was performed using the R package p*ROC*. Clusters with less than 5 cells were omitted from the statistical analysis. We consider p_adj_ ≤ 0.05 significant. (p_adj_ ≤ 0.05: *, p_adj_ ≤ 0.01: **, p_adj_ ≤ 0.001: ***, p_adj_ ≤ 0.0001: ****).

## Supporting information

Supplemental Experimental Proceedures

## Resource availability

### Lead contact

Requests for further information and resources should be directed to and will be fulfilled by the lead contact, Bjørn Tore Gjertsen, MD, PhD

### Materials availability

This study did not generate new unique reagents.

## Acknowledgments

We acknowledge the contribution from patients and thank the work by study personnel at all study sites participating in both the NordCML007 and ENEST1st clinical trials. The single-cell data was collected using the Helios and CyTOF XT mass cytometers at the Flow & Mass Cytometry Core Facility, Department of Clinical Science, University of Bergen. The Helios mass cytometer was funded by the Trond Mohn Research Foundation

## Author contributions

Conceptualization B.T.G., and S.E.G.; Data curation B.T.G., H.H.H. D.W. and S.E.G.; Formal analysis B.T.G., and S.E.G; Funding acquisition B.T.G., H.H.H. and S.E.G.; Investigation B.T.G., H.H.H., D.W., M.H., and S.E.G.; Methodology B.T.G., and S.E.G.; Project administration B.T.G., H.H.H. D.W., and S.E.G.; Resources/Provided clinical samples and data U.O.S., J.S., J.R., T.G.D., D.W., S.M., B.T.G. and H.H.H.; Software Ø.S., M.B., E.C., R.T. and S.E.G.; Supervision B.T.G. and S.P.; Validation B.T.G., E.C., R.T.; Visualization B.T.G., M.B. and S.E.G.; Writing – original draft B.T.G and S.E.G.; Writing – review & editing all authors.

