## Supplemental figures and tables for "Deep single-cell immune and signaling profiles predict long term therapy response in chronic myeloid leukemia within hours"

| Supplementary table S1 **–** Panel of antibodies used for mass cytometry | | |
| --- | --- | --- |
| Isotope | Antibody | Clone |
| 89Y | CD45 | *HI30* |
| 111Cd | CD3 | *UCHT1* |
| 112Cd | CD34 | *581* |
| 113Cd | CD123 | *6H6* |
| 114Cd | CD66b | G10F5 |
| 116Cd | HLA-DR | *L243* |
| 141Pr | CD38 | *HIT2* |
| 142Nd | cCaspase 3 | *D3E9* |
| 143Nd | pCRKL Y207 | *Polyclonal* |
| 144Nd | pTyrosine | *Tyr100* |
| 145Nd | CD4 | *RPA-T4* |
| 146Nd | CD49d | *9F10* |
| 147Sm | CD20 | *2H7* |
| 148Nd | CD16 | *3G8/B73.1* |
| 149Sm | CD25 | *2A3* |
| 150Nd | pSTAT5 Y694 | *47* |
| 151Eu | pSTAT3 S727 | *D4X3C* |
| 152Sm | CD13 | *WM15* |
| 153Eu | pSTAT1 Y701 | *58D6* |
| 154Sm | CD45RA | *Hi100* |
| 155Eu | CD27 | *L128* |
| 156Eu | p-p38 T180/Y182 | *D3F9* |
| 157Eu | CD8 | *Hit8a* |
| 158Eu | pSTAT3 Y705 | *4/P-STAT3* |
| 159Tb | pMAPKAPK2 T334 | *27B7* |
| 160Eu | CD14 | *M5E2* |
| 161Dy | CD26 | *BA5b* |
| 162Dy | FoxP3 | *PCH101* |
| 163Dy | CD56 | *NCAM16.2* |
| 164Dy | CD15 | *W6D3* |
| 165Ho | pCREB S133 | *87G3* |
| 166Er | MPO | *MPO421-BB2* |
| 167Er | IL1-RAP | *89412* |
| 168Er | CD117 | *YR145* |
| 169Tm | CD33 | *P67.6* |
| 170Er | pSRC Y418 | *SC1T2M3* |
| 171Yb | pERK1/2 T202/Y204 | *D13.14.4E* |
| 172Yb | pS6 S235/S236 | *N7-548* |
| 173Yb | STAT3total | *124H6* |
| 174Yb | CD11c | *Bu15* |
| 175Lu | CXCR4 | *12G5* |
| 176Yb | pS6 S240/S244 | *D68F8* |
| 209Bi | CD11b | *ICRF44* |

| Supplementary table S2 **–** Overview of cell surface markes used in clustering, black means used. | | | | | | | |
| --- | --- | --- | --- | --- | --- | --- | --- |
| Isotope | Antibody | Cytosplore | T cells | Mono/DC | NKs | HSPCs | Neutrophils |
| 89Y | CD45 |  |  |  |  |  |  |
| 111Cd | CD3 |  |  |  |  |  |  |
| 112Cd | CD34 |  |  |  |  |  |  |
| 113Cd | CD123 |  |  |  |  |  |  |
| 114Cd | CD66b |  |  |  |  |  |  |
| 116Cd | HLA-DR |  |  |  |  |  |  |
| 141Pr | CD38 |  |  |  |  |  |  |
| 145Nd | CD4 |  |  |  |  |  |  |
| 146Nd | CD49d |  |  |  |  |  |  |
| 147Sm | CD20 |  |  |  |  |  |  |
| 148Nd | CD16 |  |  |  |  |  |  |
| 149Sm | CD25 |  |  |  |  |  |  |
| 152Sm | CD13 |  |  |  |  |  |  |
| 154Sm | CD45RA |  |  |  |  |  |  |
| 155Eu | CD27 |  |  |  |  |  |  |
| 157Eu | CD8 |  |  |  |  |  |  |
| 160Eu | CD14 |  |  |  |  |  |  |
| 161Dy | CD26 |  |  |  |  |  |  |
| 162Dy | FoxP3 |  |  |  |  |  |  |
| 163Dy | CD56 |  |  |  |  |  |  |
| 164Dy | CD15 |  |  |  |  |  |  |
| 166Er | MPO |  |  |  |  |  |  |
| 167Er | IL1-RAP |  |  |  |  |  |  |
| 168Er | CD117 |  |  |  |  |  |  |
| 169Tm | CD33 |  |  |  |  |  |  |
| 174Yb | CD11c |  |  |  |  |  |  |
| 175Lu | CXCR4 |  |  |  |  |  |  |
| 195Pt | Cisplatin |  |  |  |  |  |  |
| 209Bi | CD11b |  |  |  |  |  |  |

### Supplemental Figures

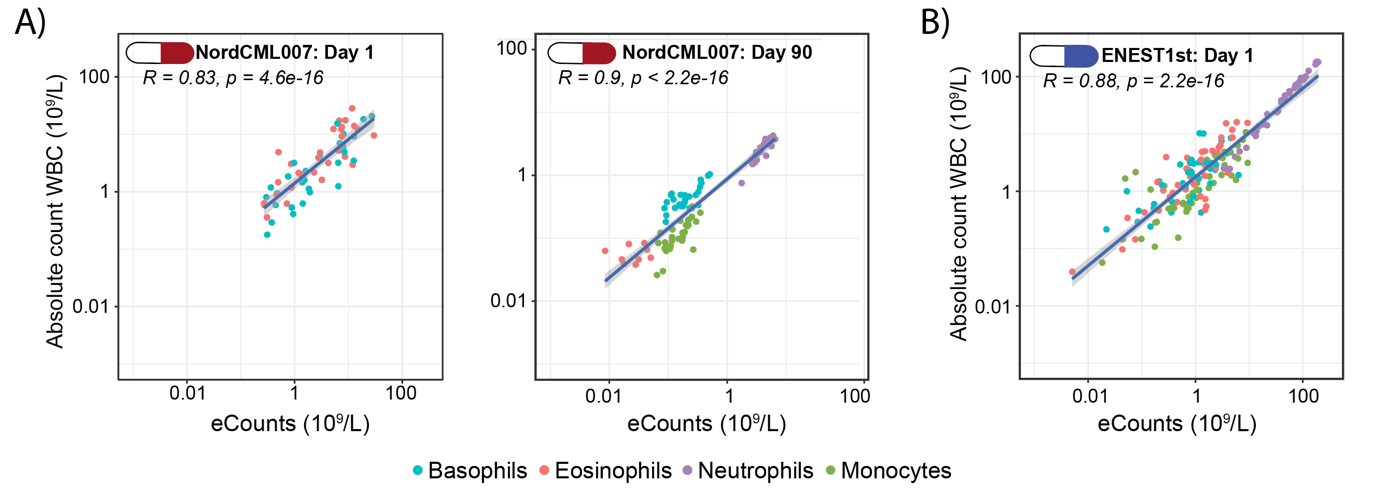

**Supplemental Figure–S1.** Spearman correlation between mass cytometry–derived eCounts and routine absolute white blood cell (WBC, 10^9^ cells/L) counts from differential white blood cell analyses in the (**A**) NordCML007 trial before dasatinib on day 1 and day 90, and in the (**B**) ENEST1st trial before nilotinib on day 1.

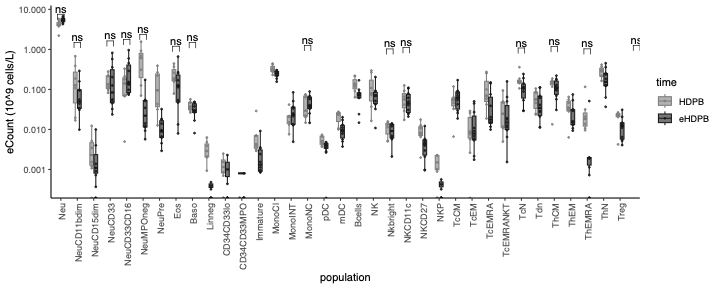

**Supplemental Figure–S2.** Comparison of eCounts from healthy donor peripheral blood samples collected alongside patient samples in the NordCML007 trial (HDPB, light grey) and ENEST1st trial (eHDPB, dark grey). All comparisons are based on grouped t-tests with for multiple testing. Significance levels are indicated as follows: Padj ≤ 0.05: *, Padj ≤ 0.01: **, Padj ≤ 0.001: ***, Padj ≤ 0.0001: ****.

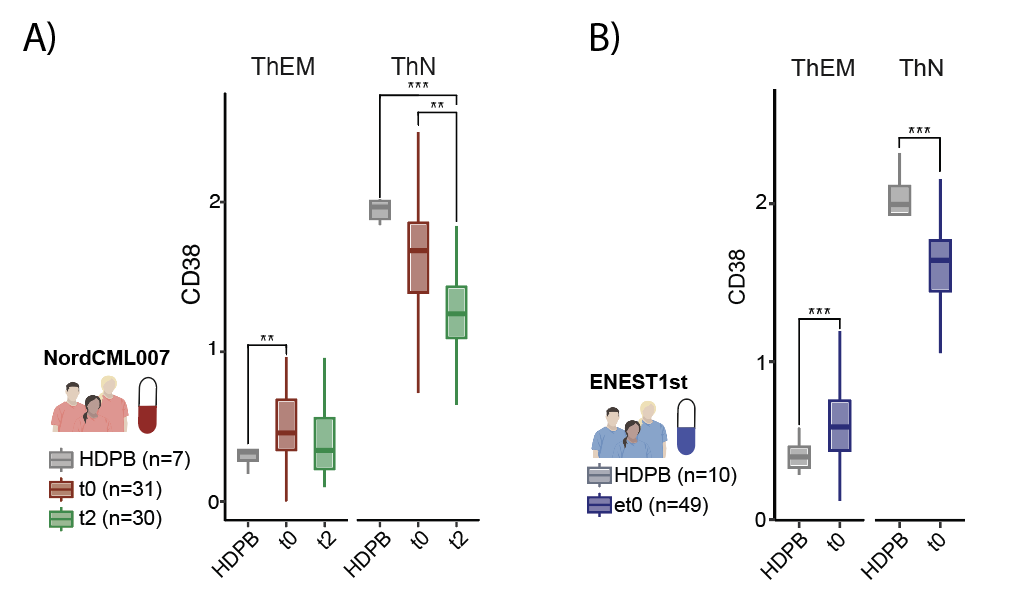

**Supplemental Figure–S3.** Boxplots showing the 75th-percentile expression of CD38 in selected T cell subsets in the (A) NordCML007 trial and (B) the ENEST1st trial. All comparisons are based on grouped t-tests with for multiple testing. Significance levels are indicated as follows: Padj ≤ 0.05: *, Padj ≤ 0.01: **, Padj ≤ 0.001: ***, Padj ≤ 0.0001: ****.

| A)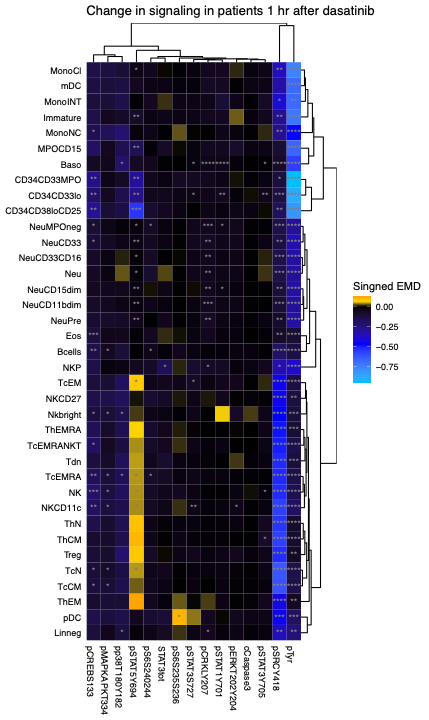 | B)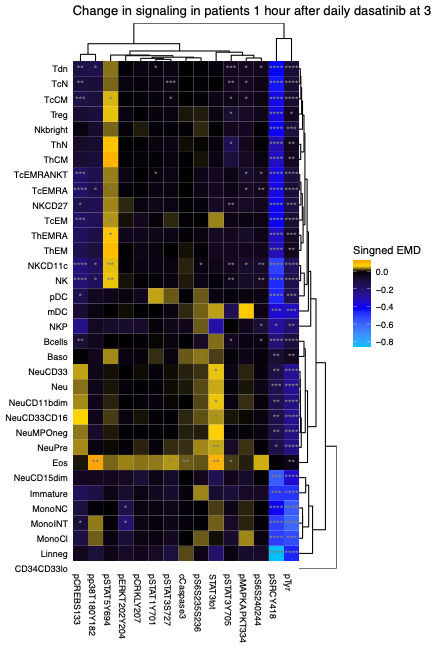 |
| --- | --- |
| C)  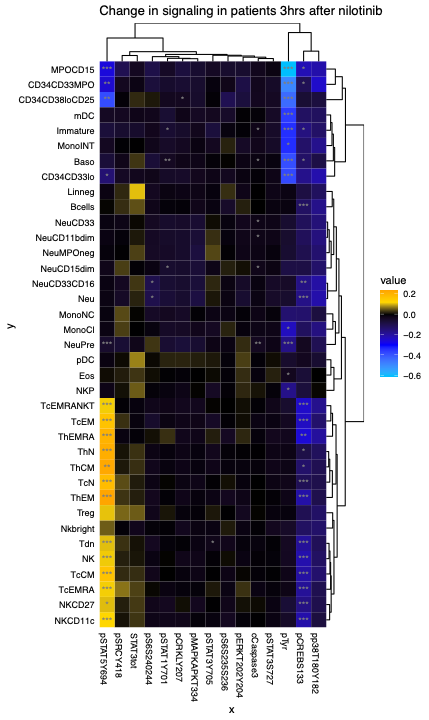 | **Supplemental Figure–S4.** Same data as in **Figure–4C and D** but shown as a heatmap of the average signed EMD comparing the change one hour after dasatinib on (A) day 1 and (B) day 90, and three hours after nilotinib on (C) day 1. The stars indicate the FDR-adjusted and paired t-test of the 75^th^ percentile of each BCR::ABL1 related signaling marker in each immune population. |

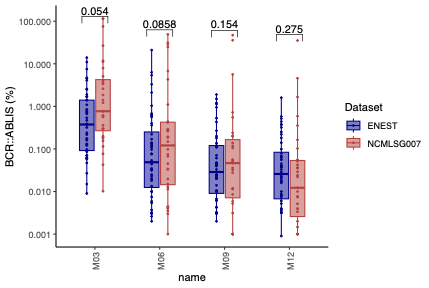

**Supplemental Figure–S5.** Grouped t-test comparing BCR::ABL^IS^ every three months the first year of treatment in the ENEST1st (blue) and NordCML007 (ref) clinical trials. Statistical comparisons were performed using unpaired t-tests with false discovery rate (FDR) correction for multiple testing. No significant differences were observed between the two cohorts for any cell population (all FDR-adjusted p > 0.05, marked as "ns").

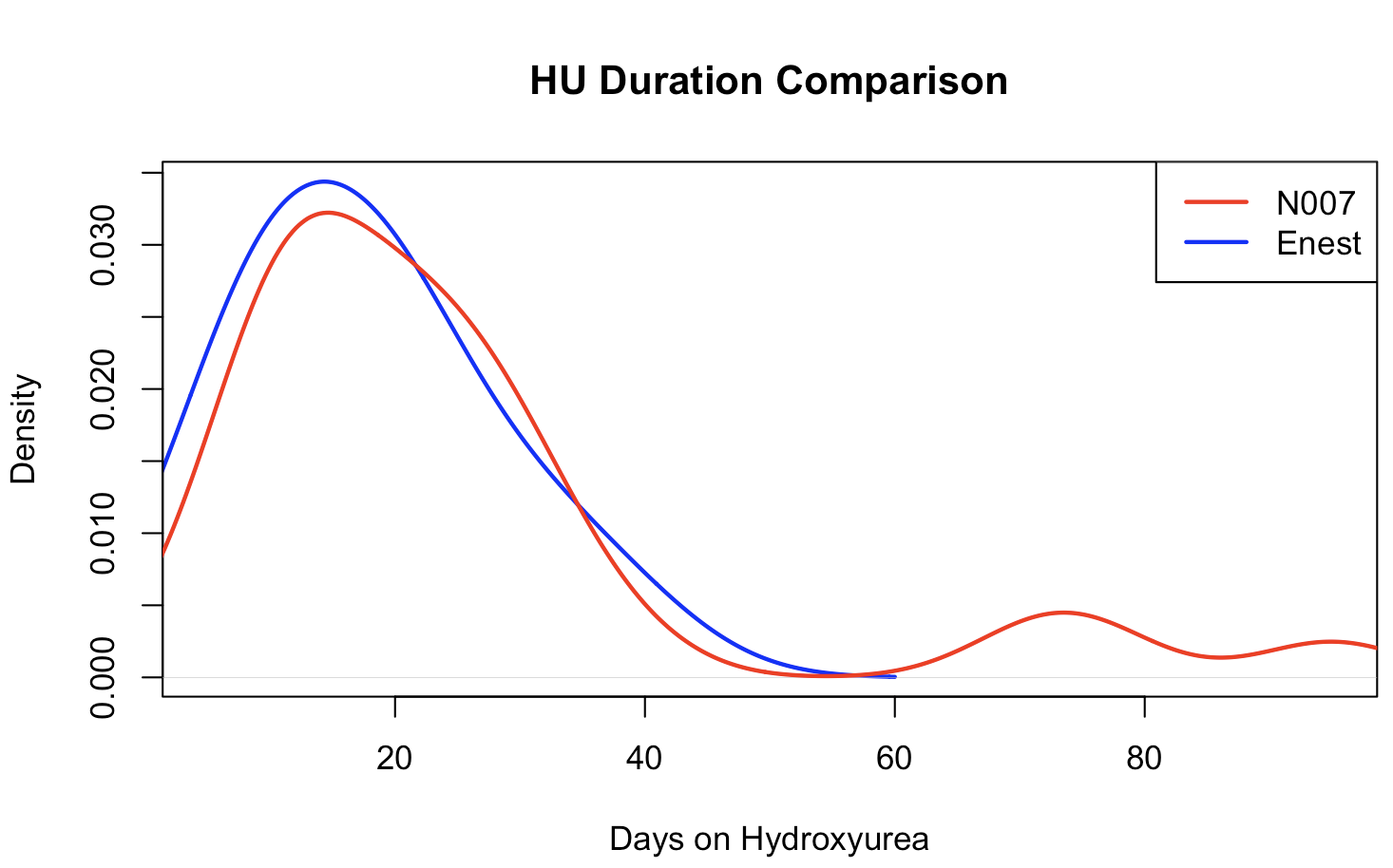

**Supplemental Figure–S6.** Kernel density estimates showing the distribution of hydroxyurea (HU) treatment duration in days for patients from the N007 study (blue line) and the ENEST cohort (red line,). Only patients with documented HU exposure (duration >0 days) were included in the analysis (NordCML007; n=19, ENEST1st; n=28). Median treatment duration was 17 days (IQR: 10-22) for N007 versus 21 days (IQR: 11.75-28.50) for ENEST. The distributions did not differ significantly between cohorts (Mann-Whitney U test, W = 214, p = 0.26). Bandwidth for density estimation was set to 7 days for the N007 cohort.
