## Supplemental Experimental Proceedures for "Deep single-cell immune and signaling profiles predict long term therapy response in chronic myeloid leukemia within hours"

Supplemental Experimental Procedures

### 1. Identification of eosinophils by increased uptake of cisplatin

Identification of eosinophils in whole blood samples following preservation by formaldehyde containing buffers (e.g. BD Lyse/Fix) is challenging. The mass cytometry platform naturally lacks the ability to measure the high side-scatter (SSC) of eosinophils. Many of the cell surface antigens that can be used to distinguish eosinophils from other immune cells are not reliably detected after preservation with formaldehyde.(1) One defining feature of eosinophils are their cytoplasmic granules that contain high levels of arginine-rich cationic proteins. If not blocked with heparin, these cationic proteins will attract the net-anionic chelator-metal complex conjugated to antibodies, causing unspecific binding.(2)

During barcoding of Lyse/Fix preserved whole blood samples with cisplatin isotopes we noticed a recurring population of cells with increased Pt levels. The phenotype of these cells was granulocytic (CD45^low^CD66b^+^CD15^-^CD123^-^), but not neutrophils nor basophils (**Supplemental Experimental Procedures Figure S1**). We hypothesized that this population was indeed eosinophils, and that the increased Pt labelling was the result of an effect like that seen with the chelator-metal complex conjugated to antibodies.

We devised a barcoding scheme where samples were intially barcoded and pooled using an in-house formulation of the 6-chooses-3 palladium 20-plex metal barcoding developed by Zunder.(3) Then, two 20-plex pools were barcoded and pooled using two unique combinations of two cisplatin isotopes (4-chooses-2, **Supplemental Experimental Procedures Table 1**).(4) The two pools were separated based in ^194^Pt and ^198^Pt, while all cells were equally stained with ^195^Pt for use in identification of eosinophils.

| **Supplemental experimental procedures Table 1:** Cisplatin was used to barcode and pool of two pools of cells already barcoded and pooled using the 20-plex Palladium barcoding reagent. All cells were labelled with Pt^195^ | | | |
| --- | --- | --- | --- |
|  | Pt^194^ | Pt^195^ | Pt^198^ |
| 20-plex pool #1 | x | x |  |
| 20-plex pool #2 |  | x | x |

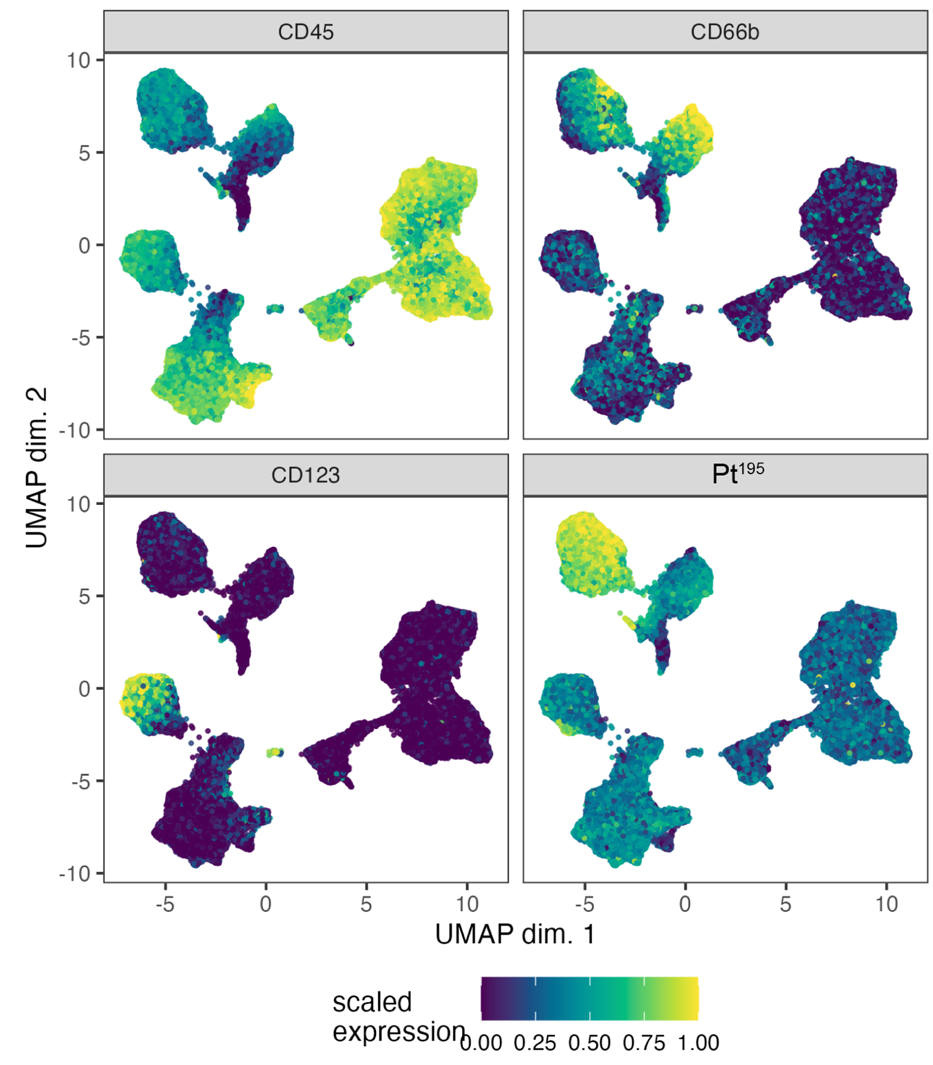

**Supplemental experimental procedures-S1 – Detection of Eosinophils.** UMAP analysis of single cell data as in **Figure 1-B and C**, but colored cell surface marker expression to visualize utility of Pt^195^ in detection of Eosinophils

### 2. Calculation of absolute counts of mass cytometry immune populations (eCounts)

Routine white blood cell absolute counts (WBC, 10⁹ cells/L) were obtained at study visits in the NordCML007 and ENEST1st clinical trials. Measurements were performed on the same day as mass cytometry sample collection, corresponding to timepoints before treatment initiation (t0/et0) and after three months of dasatinib treatment (t2). The absolute count of the immune populations identified in the mass cytometry data (eCounts) was calculated by dividing the proportion (%) of each population with 100 and then multiplying with the WBC.

eCounts = $\frac{Proportion (\%)}{100\%}$ × WBC

For the samples collected 1 hour after per oral dasatinib (t1 and t3), corresponding routine WBC measurements was obtained limited number of patients at day 90 (none at day 1). To be able to generate eCounts from these post-TKI samples, we first evaluated if the WBC from before dasatinib could be used to calculate the eCounts 1 hour after dasatinib. Comparing these eCounts of the immune populations before and 1 after dasatinib after 90 days, we observed a statistically significant reduction in eosinophils (p_adj_=1.19x10^-4^) (**Supplemental Experimental Procedures Figure S2-A**). This is not in line with Mustjoki and colleagues report (5) where the eosinophil count was reported to be largely unaffected by dasatinib after 1-2 hours. Mobilization of leukocytes into the blood stream after dasatinib results in a significantly increased WBC, likely causing this artefact. Indeed, not adjusting for an increased WBC would result in an apparent decrease in eosinophils, and, importantly, an underestimated eCounts of all immune populations in general. However, during sample preparation, we extracted the exact same volume of sample volume from samples collected before and after dasatinib, capturing any changes in cellular concentration (mobilization) after 1 hour. This enabled us to estimate the WBC after dasatinib by adjusting the WBC before using the captured mobilization.

To test this estimation, we first compared the WBC from the limited number of patients where we had ground truth WBC 1 h after dasatinib measurements, with the corresponding WBC measured before dasatinib (**Supplemental Experimental Procedures Figure S2-B**). We found that although they were similar, these two measurements were not significantly correlated, showing that the dasatinib induced mobilization affected the WBC one hour after per-oral dosing. We then compared our estimated WBC 1 hour after dasatinib with ground truth WBC and found that good agreement between our estimated value and the true values. This approach was applied for samples collected on day 1 (t0/t1) and after three months of dasatinib (t2/t3). We did not adjust the WBC values for patients treated with nilotinib.

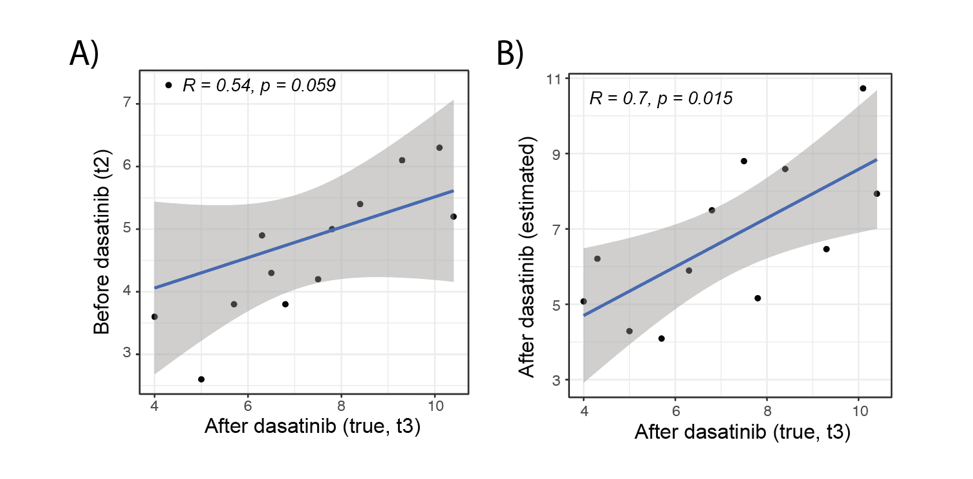

**Supplemental Experimental Procedures Figure S2 – (A)** Comparison of estimated absolute counts (eCounts) of immune cell populations before (t2) and 1 hour after (t3) dasatinib administration at day 90, calculated using pre-dasatinib WBC values. **(B)** Validation of estimated post-dasatinib WBC values. Estimated WBC values at 1 hour post-dasatinib, calculated by adjusting pre-dasatinib WBC using the mobilization captured during sample processing.
